# Evolution of NELL binding by dual-ligand-responsive axon guidance receptor Robo

**DOI:** 10.64898/2026.09.23.753883

**Authors:** Joseph S. Pak, Lakshmi Prakash, Yeonwoo Park, Wioletta I. Nawrocka, Raunak Kundagrami, Joseph W. Thornton, Alexander Jaworski, Engin Özkan

**Affiliations:** Department of Biochemistry and Molecular Biology, The University of Chicago, Chicago, Illinois, USA; Institute for Neuroscience, The University of Chicago, Chicago, Illinois, USA; Institute for Biophysical Dynamics, The University of Chicago, Chicago, Illinois, USA; Department of Neuroscience, Brown University, Providence, Rhode Island, USA; Robert J. and Nancy D. Carney Institute for Brain Science, Brown University, Providence, Rhode Island, USA; Committee on Genetics, Genomics, and Systems Biology, The University of Chicago, Chicago, Illinois, USA; Department of Human Genetics, The University of Chicago, Chicago, Illinois, USA; Department of Ecology and Evolution, The University of Chicago, Chicago, Illinois, USA

**Keywords:** Robo receptors, NELL ligands, axon guidance, neural development, Slit signaling, chordate evolution, gene duplication, subfunctionalization, liquid–liquid phase separation (LLPS), receptor–ligand interactions

## Abstract

Robo receptors are conserved across bilaterians and best known for their ability to mediate axonal repulsion in response to Slit family ligands. In mammals, this applies to Robo1 and Robo2, but mammalian Robo3 binds NELL proteins instead of Slits. The evolutionary origin of NELL-Robo interactions and the possible existence of dual-ligand responsiveness across species remain unknown. Here, we systematically analyzed Robo and NELL homologs across bilaterians and found that NELL-Robo binding is conserved among chordate Robos, but not in protostomes. We show that cephalochordate Robo and NELL can mediate axon repulsion *in vitro*, suggesting conserved functionality. We observed that conformational masking of the NELL-binding site is prevalent among chordate Robos, modulating NELL-Robo affinity. We also demonstrate that NELL–Robo complexes undergo liquid–liquid phase separation *in vitro*, a property preserved from cephalochordates to mammals. Our findings support a model in which an ancestral chordate Robo receptor was dual-responsive to Slit and NELL, still the case for some extant Robos, and vertebrate paralogs subfunctionalized, with full ligand specialization emerging in mammals.

## INTRODUCTION

Establishing precise neuronal connectivity is one of the most complex and tightly regulated processes in animal development. Axon guidance, in which growing axons interpret extracellular cues to navigate towards appropriate targets, is a critical step in neural circuit assembly (Kolodkin & Tessier-Lavigne, 2011). Among the best-characterized systems governing axon pathfinding is the Slit-Robo signaling axis, which mediates repulsion to aid in the assembly of various neuronal circuits. Robo (Roundabout) receptors are conserved across bilaterians and are expressed on axonal growth cones, where they signal in response to secreted Slit ligands. All three neuronal Robo paralogs in mammals (Robo1-3) and all three Slits (Slit1-3) are expressed in the developing spinal cord, where they gate axonal crossing of the ventral midline (Dickson & Gilestro, 2006). Loss of Robo or Slit function leads to midline crossing defects in mice, zebrafish, fruit flies, and nematodes, underscoring the evolutionary conservation and developmental importance of this signaling system (Kidd *et al*, 1998a, 1999; Brose *et al*, 1999; Evans & Bashaw, 2010). Furthermore, Robos have been demonstrated to be Slit receptors across bilaterian model organisms (Kidd *et al*, 1999; Brose *et al*, 1999; Yuan *et al*, 1999; Zelina *et al*, 2014), and Slit-Robo signaling fulfills various critical functions beyond axon guidance, contributing to developmental processes as diverse as angiogenesis and mammary gland morphogenesis (Blockus & Chédotal, 2016).

While Slit-Robo signaling is broadly conserved, important mechanistic differences have emerged across species. For example, *Drosophila* and vertebrate Robos are internalized by unrelated E3 ubiquitin ligase adaptors (Gorla *et al*, 2019; Kidd *et al*, 1998b; Keleman *et al*, 2002). The most striking divergence is found in mammalian Robo3, which has lost its ability to bind Slits (Zelina *et al*, 2014; Mambetisaeva *et al*, 2005), yet is still essential for midline crossing (Marillat *et al*, 2004; Sabatier *et al*, 2004; Di Meglio *et al*, 2008; Jaworski *et al*, 2010). Recent work has shown that mammalian Robo3 instead binds NELL1 and NELL2, a structurally distinct family of secreted ligands that also induce axonal repulsion (Jaworski *et al*, 2015; Pak *et al*, 2020). These interactions occur through a unique site on the Robo3 ectodomain (ECD), the first Fibronectin type-III (FN) domain (Jaworski *et al*, 2015; Pak *et al*, 2020), separate from the canonical Slit-binding interface on Robo1 and Robo2, the first immunoglobulin (IG) domain (Chen *et al*, 2001; Liu *et al*, 2004; Morlot *et al*, 2007) (**Figure 1A**). Thus, Robo3 appears to have acquired a different ligand while retaining its role in commissural axon guidance, indicating that Robo receptors are more functionally diverse than previously thought.

**Figure 1.**
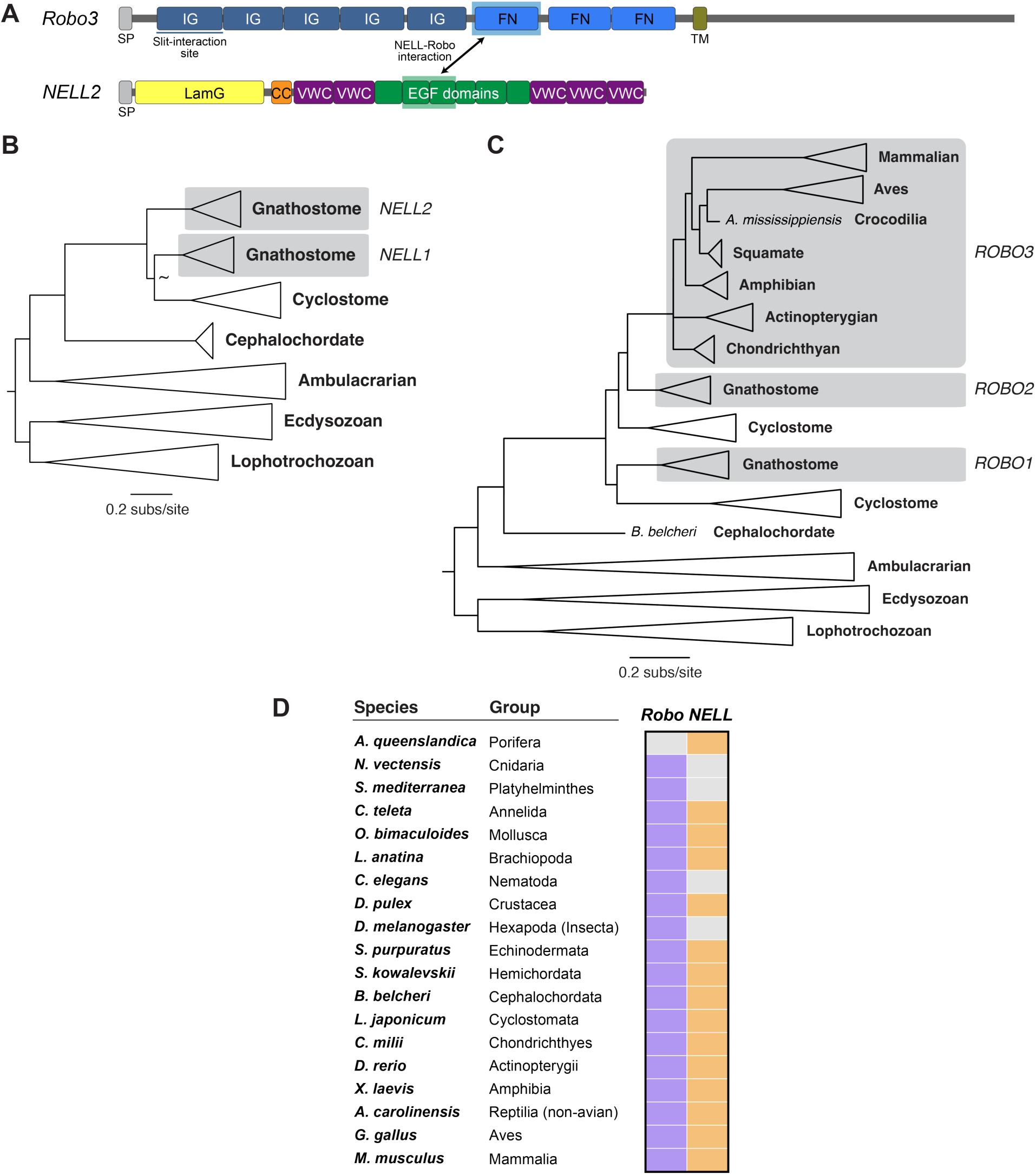
Distribution of NELL and Robo genes across metazoans. **A.** Domain architecture of NELL and Robo proteins, labeling conserved immunoglobulin (IG), Fibronectin type-III (FN), Laminin G (LamG), Epidermal Growth Factor (EGF) and von Willebrand Factor type-C (VWC) domains, signal peptides (SP), a transmembrane (TM) helix and a coiled coil (CC). The distinct and non-overlapping Slit and NELL-interaction sites on Robos are highlighted. **B,C.** Maximum-likelihood phylogenetic tree of NELL and Robo orthologs from representative metazoan clades. **D.** Major metazoan lineages annotated for presence or absence (gray) of NELL and Robo genes.

Recent biochermical analysis has shed some light on how mammalian Robos have subfunctionalized to use separate ligands: mammalian Robo1 and Robo2 possess cryptic NELL binding, likely due to hairpin formation of the Robo1/2 ECD and blocking of the NELL-binding site (Pak *et al*, 2020; Yamamoto *et al*, 2019; Miyaguchi *et al*, 2022; Barak *et al*, 2019; Aleksandrova *et al*, 2018). Even when the conformational block is removed, the NELL–Robo1 interaction is weak (Pak *et al*, 2020). Mammalian Robo3, on the other hand, has key Slit-interacting amino acids mutated, abolishing Slit binding (Zelina *et al*, 2014). However, how ligand-Robo affinities evolved in non-mammalian vertebrates remains a mostly open question (Yu *et al*, 2014).

Despite growing evidence for NELL functions in mammals, the evolutionary origin of NELL-Robo interactions remains poorly understood. Four Robos (including the non-neuronal Robo4, which has lost several domains) in vertebrates arose through two rounds of duplication that occurred after the divergence of cephalochordates (Xu *et al*, 2021). It is unclear whether NELL responsiveness is a mammalian innovation or reflects an ancestral feature of Robo receptors that was retained in Robo3 but lost in Robo1 and Robo2. In this context, understanding whether Robo paralogs subfunctionalized by partitioning ancestral Slit and NELL responsiveness, or whether NELL binding represents a neofunctionalization of Robo3, is critical to reconstructing the functional evolution of the Robo family.

To address these questions, we performed a comprehensive phylogenetic, biochemical, and functional analysis of Robo and NELL homologs across bilaterians. We found that NELL binding is broadly conserved among deuterostome Robos, including non-mammalian Robo1, Robo2, and Robo3 orthologs, but is absent in protostome Robos, which may only bind Slits. We further demonstrated that the NELL-Robo signaling axis is functional in basal chordates: the lancelet NELL-Robo pair is sufficient to mediate repulsion in a mammalian axon turning assay, indicating conserved guidance activity. Similarly, zebrafish Robo1 and Robo3 respond to NELL in this assay, further supporting chordate Robo1 as functional NELL receptors outside mammals. In addition, we observed that NELL–Robo complexes undergo liquid–liquid phase separation (LLPS) *in vitro*, a biophysical property that is conserved from cephalochordates to mammals.

Together, our findings support a model in which the ancestral chordate Robo receptor was dual-responsive to Slit and NELL ligands, and this dual functionality was later partitioned among Robo paralogs following genome duplication. This subfunctionalization culminated in the complete division of ligand specificities observed in mammals, with Robo1 and Robo2 mediating Slit-dependent guidance and Robo3 adopting a specialized role in NELL-mediated signaling, highlighting the evolutionary depth and functional diversification of the Robo family.

## RESULTS

### NELL family members are conserved across metazoans

NELL proteins were originally characterized as ligands for mammalian Robo3, but their distribution outside vertebrates has not been systematically evaluated. To address this gap, we performed a comprehensive homology search across publicly available genomes and transcriptomes. We observed that NELLs retain their domain topology throughout metazoa (**Figures 1A, S1A**).

Our search recovered NELL orthologs across major deuterostome lineages, including vertebrates, lancelets (cephalochordates), echinoderms, and hemichordates (**Figure 1B**). Strikingly, we also identified NELL genes in several protostome clades, most notably within Spiralia (mollusks, annelids, and brachiopods) and in a subset of arthropods, including horseshoe crabs and several crustaceans (**Figure 1B**).

Conversely, we found no trace of NELL sequences in the intensively studied model organisms *Drosophila melanogaster* and *Caenorhabditis elegans*, nor in any other surveyed nematode genomes, suggesting lineage-specific losses of the gene in these taxa. The presence of NELLs in basal protostomes, coupled with their absence in select derived lineages, implies multiple independent loss events rather than a late origin within deuterostomes.

Unexpectedly, we also identified a NELL-like sequence in a sponge (*Amphimedon queeslandica*), which implies that the NELL family predates the origin of bilaterian animals and their nervous system. Although divergent, the *A. queenslandica* protein retains the characteristic domain architecture of NELLs and the typical cysteine pattern of EGF and VWC domains (**Figure S1**), and clusters with chordate NELLs in phylogenetic analyses.

In contrast, Robo receptors have been detected across Bilateria and in several cnidarians, such as jellyfish and sea anemones, but not in sponges (Heger *et al*, 2020). The presence of a NELL-like gene in *A. queenslandica* without a corresponding Robo ortholog supports the idea that canonical NELL–Robo signaling evolved later, likely in early eumetazoans, and was subsequently elaborated in chordates (**Figures 1C, D**).

Together, these data demonstrate that NELL family members are far more ancient and taxonomically widespread than previously appreciated, spanning the full breadth of deuterostomes and extending deep into protostomes and even non-bilaterian animals such as sponges (**Figures 1B,D**). The patchy distribution in protostomes highlights dynamic evolutionary trajectories, while the pervasive presence in chordates sets the stage for investigating how NELL– Robo signaling diversified alongside vertebrate nervous system complexity.

### NELL binding is a conserved feature among chordate Robo orthologs

To determine when NELL binding evolved among Robos, we constructed a maximum-likelihood phylogeny of bilaterian Robo ectodomains (**Figure 1C**). Eight species spanning the bilaterian diversity were selected for biochemical interrogation: three protostomes—mosquito (*Aedes aegypti*), scorpion (*Centruroides sculpturatus*), and octopus (*Octopus bimaculoides*)—and five deutrostomes—lancelet (*Branchiostoma belcheri*), lamprey (*Lethenteron japonicum*), zebrafish, frog, and chicken.

To study if the Robo and NELL orthologs we identified bind each other, we codon-optimized, synthesized, and expressed in insect cells constructs comprising NELL EGF domains 1–3 and Robo’s first FN domains (FN1) as bait (Fc-tagged) or prey (Alkaline phosphatase-tagged), as these domains are known to fully mediate the NELL–Robo interaction (Pak *et al*, 2020), and tested NELL-Robo binding using a high-throughput pairwise interaction assay, the extracellular interactome assay (ECIA) (**Figure 2A**) (Nawrocka *et al*, 2026). We detected no interaction for any protostome pair. By contrast, Robo binding was observed for all deuterostome NELLs tested, including the single lancelet (*B. belcheri*) and lamprey (*L. japonicum*) NELL-Robo pairs (**Figure 2B**). These data demonstrate that NELL recognition is an ancestral feature of chordate Robos (**Figure 2C**). Together with existing data showing Slit binding across chordate Robos, our work suggests that the last common chordate ancestor possessed a dual-ligand, Slit/NELL-responsive Robo.

**Figure 2.**
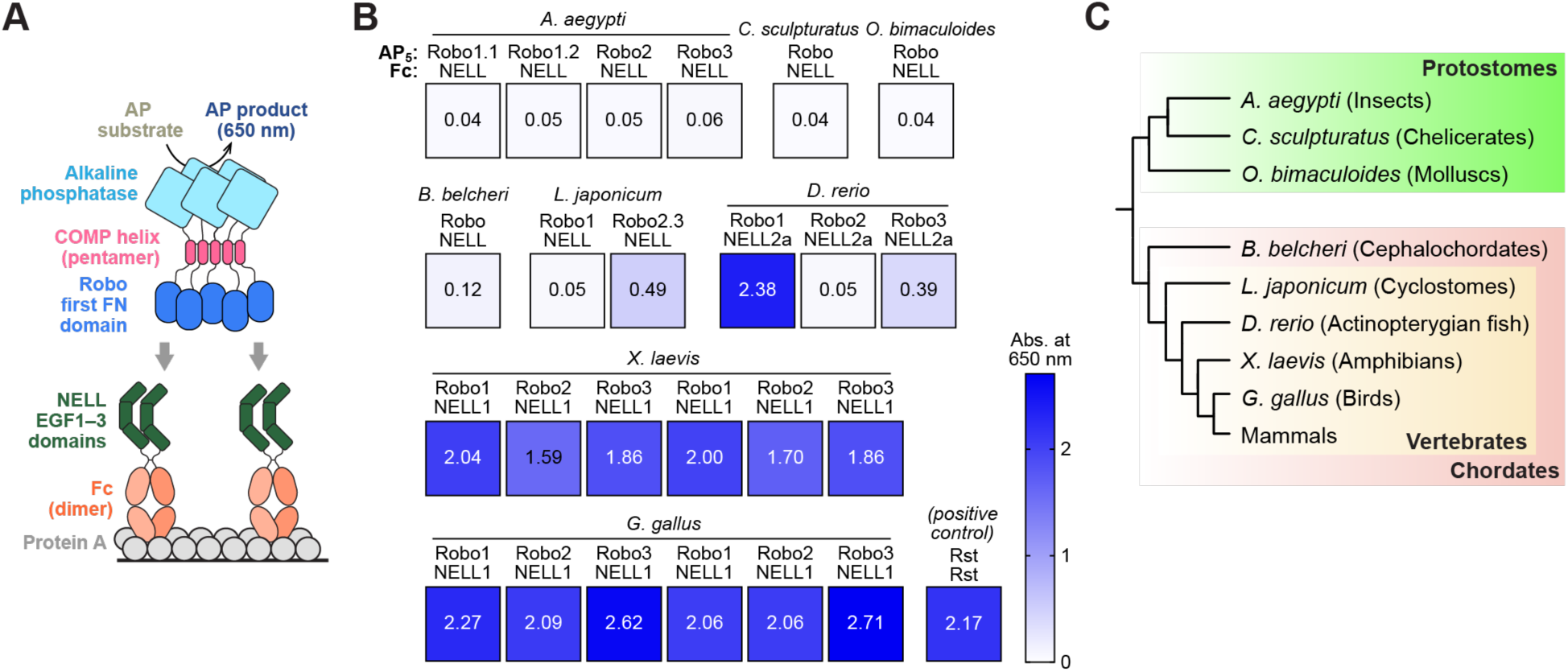
NELL-Robo interactions are conserved across deuterostomes but absent in protostomes. **A.** Schematic of the extracellular interactome assay (ECIA) in which Fc-tagged NELL serve as bait and pentamerized AP-tagged Robos as prey. **B.** ECIA results with truncated constructs (NELL EGF1-3 and Robo FN1) show that chordate Robo and NELLs have affinity towards each other, while no binding is observed for protostome NELL–Robo pairs. **C.** Phylogenetic relationships of the Robo and NELL orthologs tested for binding.

### Chordate Robo and NELLs interact with high affinity

NELL-Robo interaction has been observed to be lower affinity, or cryptic, in the context of the full Robo ECD, as the folding of the ECD can mask the NELL binding site (Pak *et al*, 2020; Yamamoto *et al*, 2019). We therefore proceeded to surface plasmon resonance (SPR) analysis to measure dissociation constants (*K*_D_) and confirm whether conformational masking, or autoinhibition, may affect native NELL–Robo interaction strength.

To measure NELL–Robo binding quantitively for various non-mammalian orthologs, we biotinylated NELLs and captured them on streptavidin-coated SPR chips. Putative cognate Robos, as complete ECD or the isolated FN1 domain, were flowed as analytes (**Figure 3**; summarized in panel **F**). For chicken proteins, we observed strong binding between Robo FN1 domains and NELLs (**Figure 3A**); however, the binding responses were significantly lower for Robo2 ECD and very low for Robo1 ECD, mimicking previous observations with mammalian NELLs and Robos (**Figure 3B**) (Pak *et al*, 2020; Yamamoto *et al*, 2019). These results demonstrate that avian Robo1 and Robo2 ECDs likely fold to occlude NELL-binding sites available in truncated FN1-only constructs, while Robo3 may strongly prefer an open state, similar to to the case in mammals. For zebrafish proteins, NELL2b interacts strongly with Robo1 and Robo3 FN1 and ECDs, while Robo2 ECD showed significantly weaker NELL2b binding compared to the high-affinity Robo2 FN1 binding to NELL2b (**Figures 3C,D**).

**Figure 3.**
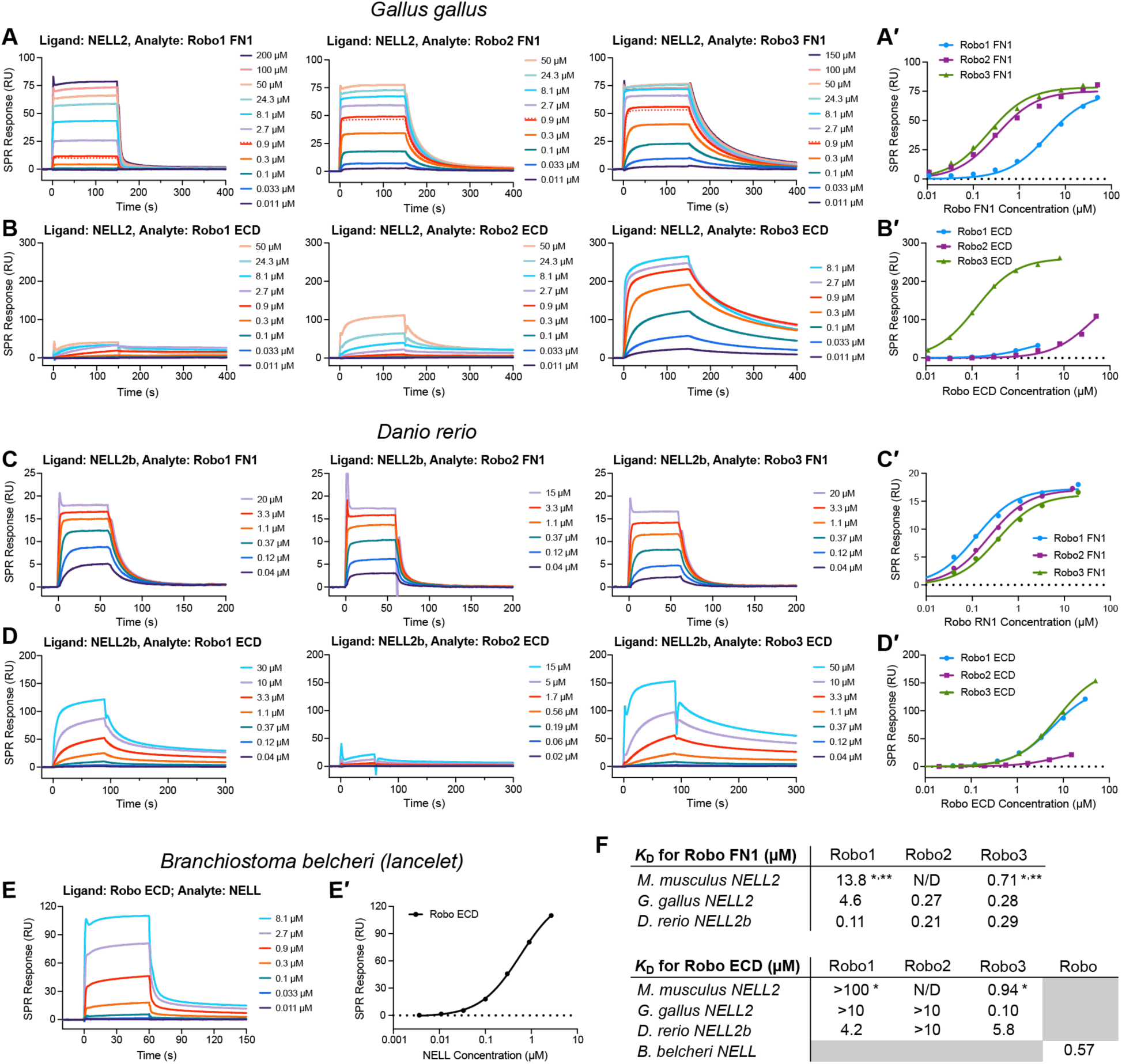
NELL–Robo interactions are high-affinity in cephalochordates and enhanced by domain truncation in vertebrates. **A,B.** SPR sensorgrams for binding between *G. gallus* Robos and full-length NELL2, using Robo FN1 domains (A) or ectodomains (B). Respective binding isotherms are plotted in A′ and B′. **C–D.** SPR sensorgrams for binding between *D. rerio* Robos and full-length NELL2b, using Robo FN1 domains (C) or ectodomains (D). Respective binding isotherms are plotted in C′ and D′. **E.** SPR sensorgram for binding between *B. belcheri* Robo ECD and full-length NELL. Binding isotherm is included in E′. **F.** Summary of measured dissociation constants. * Mouse NELL vs. Robo results are from Pak et al;. 2020. ** From mouse NELL2 EGF1-6 (ligand) against Robo FN1-3 (analyte) experiments.

In all tested Robo3 orthologs, except chicken and previously mammalian (Pak *et al*, 2020), we observed marked decrease in NELL binding when entire Robo ECDs were included (**Figs 3A′** vs. **3B′** and **3C′** vs. **3D′**). This suggests that confirmational masking of the NELL binding site in complete Robo ECDs is conserved throughout multiple vertebrate Robo lineages.

Since we have observed that the three neuronal vertebrate Robo orthologs can bind NELLs, we decided to test NELL–Robo affinity in nonvertebrate chordates, which diverged from the vertebrate line before the Robo and NELL genes expanded via duplication. The sole Robo and NELL from *Branchiostoma belcheri* interact with each other with a *K*_D_ of 0.6 µM when tested with the complete Robo ECD and full-length NELL (**Figure 3E**). This provides further evidence that the NELL-Robo interaction goes back at least to the ancestral chordates, and that Robo was originally a dual NELL and Slit receptor before it duplicated and diverged in its ligand specificity in the vertebrate line.

### NELL–Robo complexes undergo conserved phase separation *in vitro*

During initial co-purification trials, we observed that mixing NELL2 and the human Robo3 ECD consistently form an insoluble phase, often yielding gel-like pellets over time. Differential interference contrast (DIC) microscopy of clarified mixtures revealed spherical micrometer-sized condensates that emerged at protein concentrations close to the SPR-determined affinities. This phenomenon was reproducible with mouse NELL2 and Robo3, suggesting it is an intrinsic property of the complex (**Figure 4A**). SDS-PAGE of the pelleted material demonstrated a roughly 1:1 stoichiometry of Robo3 and NELL2 (**Figure 4B**). To verify that both partners occupy the same dense phase, we labelled Robo3 ECD and NELL2 at their N-termini with spectrally distinct NHS-ester fluorophores. Confocal imaging showed complete co-localization of Robo and NELL within droplets (**Figure 4C**). Overnight incubation drove the droplets to coalesce into a transparent, viscoelastic gel, indicative of liquid-to-solid maturation.

**Figure 4.**
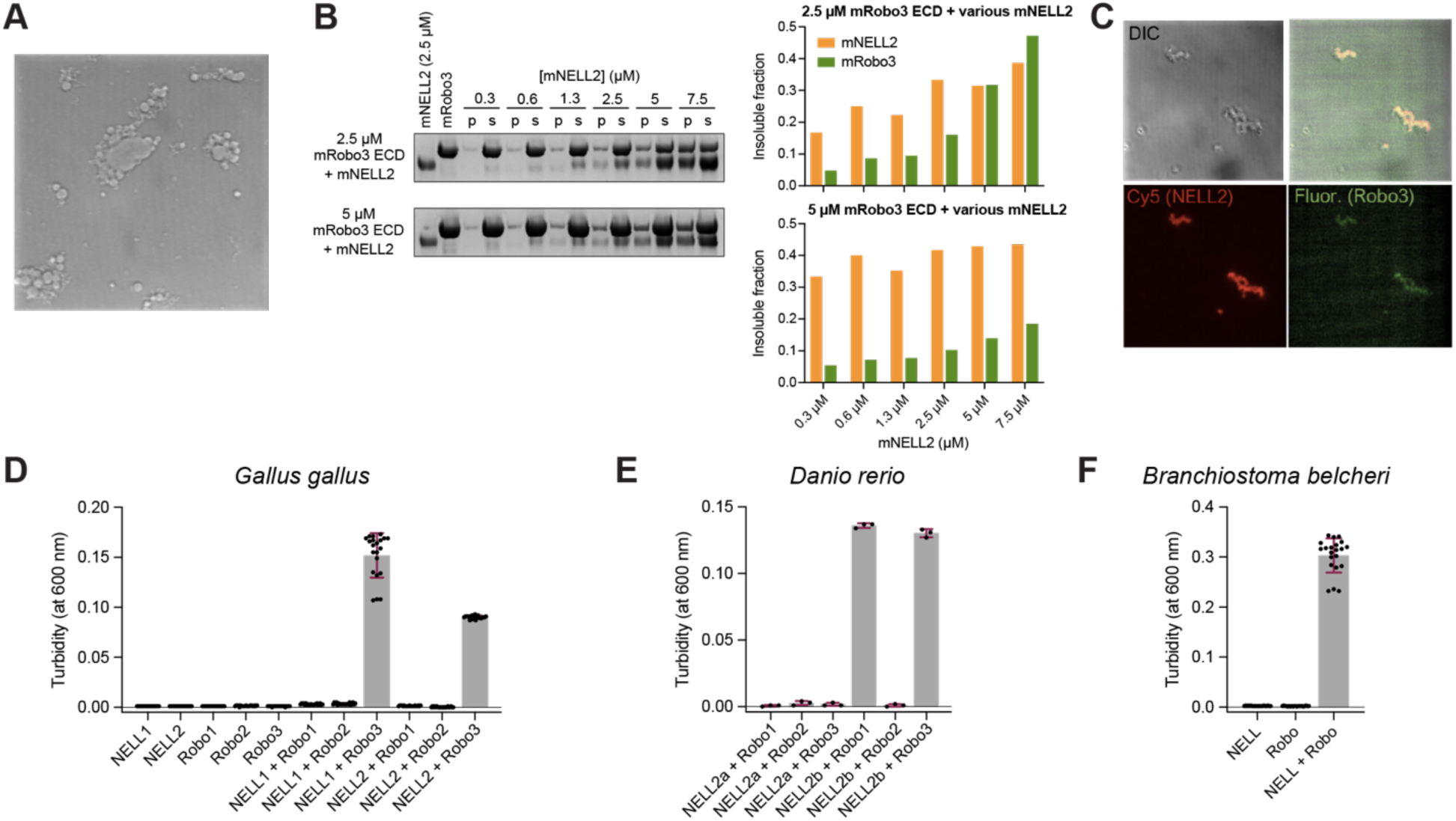
NELL–Robo complex formation drives phase separation from cephalochordate to mammals. **A.** DIC micrograph showing condensate formation upon mixing Robo3 ectodomain (14 µM) and NELL2 (5 µM) 20 minutes after mixing. **B.** SDS-PAGE of pelleted condensates showing approximately 1:1 Robo:NELL stoichiometry. **C.** Fluorescence microscopy of N-terminally labeled fluorescein-Robo3 and Cy5-NELL2 (mouse) demonstrating co-localization within condensates (40 min post-mixing). **D–F.** Conservation of phase-separation behavior across chick, zebrafish, and lancelet NELL– Robo complexes as observed by turbidity of Robo+NELL solutions measured at 600 nm.

We next asked whether condensate formation is conserved across the chordate lineage. We mixed orthologous NELLs and Robo ectodomains from the chick, zebrafish and cephalochordate *Branchiostoma belcheri*. We observed turbidity in these solutions, but only for constructs that exhibited high-affinity interactions in SPR experiments (**Figures 4D–F**). Thus, the propensity of NELL–Robo complexes to undergo concentration-dependent condensation *in vitro* appears to be a conserved molecular property of the complex among chordate orthologs.

### Cephalochordate NELL drives Robo-mediated axon guidance

Having established that cephalochordate Robo and NELL form high-affinity complexes, we next asked whether these interactions are functionally conserved in axon guidance. To test this, we employed a genetic rescue approach in the Dunn chamber axon turning assay, which was previously used to study the function of the mammalian NELL2-Robo3 complex. As reported before (Pak *et al*, 2020), embryonic spinal cord neurons from *Robo3^-/-^* mice fail to be repelled by NELL2 *in vitro*, but restoring mouse Robo3 expression via electroporation of a rescue construct restores NELL2-induced axon turning in commissural neurons; further, it confers NELL2 sensitivity to ipsilaterally projecting neurons, which normally do not respond to NELL2 at all (**Figures 5A,B,F** and **Figure S3A**). The same rescue platform was used to investigate the axon guidance activities of cephalochordate and zebrafish NELLs and Robos. We transfected embryonic *Robo3^−/−^* spinal cord neurons with the sole lancelet (*B. belcheri*) Robo, representing the ancestral state of Robos prior to gene duplications in the chordate lineage, and applied a gradient of lancelet NELL to the culture. We observed robust repulsive axon turning responses, with magnitudes comparable to those elicited by mouse NELL2-Robo3 pairs (**Figures 5C,F** and **Figure S3A**). Similarly, zebrafish Robo1- or Robo3-expressing neurons responded to zebrafish NELL2b (**Figures 5D,E,F** and **Figure S3A**). Hence, the lancelet and zebrafish NELL–Robo complexes can mediate axon repulsion, just as the mouse NELL2-Robo3 complex. As the lancelet and zebrafish proteins signal in mouse cells, these results also suggest that the downstream signaling machinery required for NELL-induced Robo repulsion is functionally conserved between lancelet, zebrafish, and mouse, supporting the idea that this ligand–receptor module was already active in the common chordate ancestor (**Figure 5G**).

**Figure 5.**
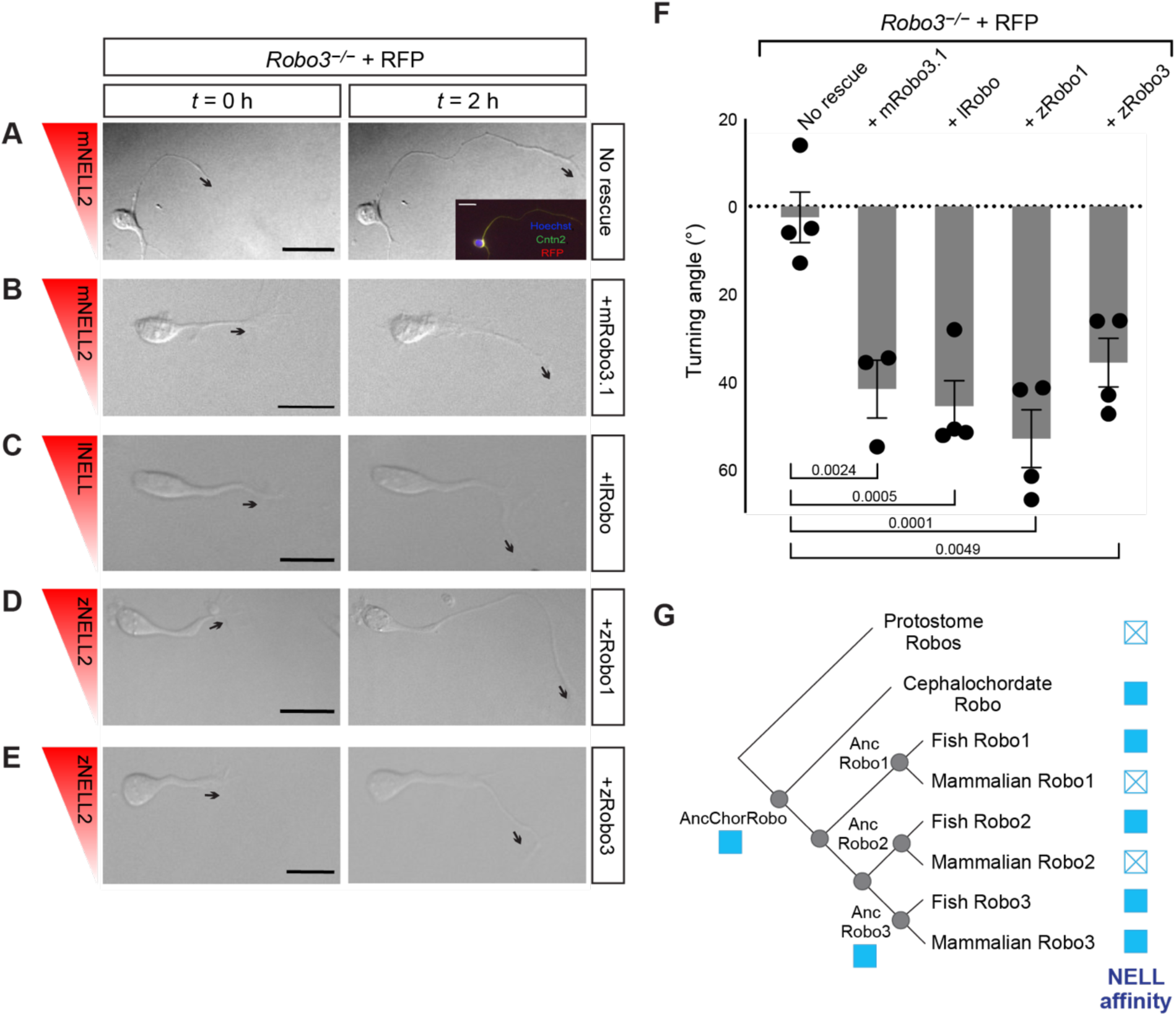
Cephalochordate and zebrafish Robos mediate axon repulsion by cognate NELLs in mouse spinal neurons. **A–E.** Representative images of *Robo3^−/−^* mouse commissural neurons transfected with various Robo expression constructs, live-imaged in 50 ng/ml NELL gradients (0-2 h). Arrows indicate orientation of distal axon segment used for measuring turning angles. Inset shows immunohistochemistry for Cntn2 and RFP, identifying the imaged neuron as commissural and having received the RFP mock rescue construct. Scale bars: 20 μm. **F.** Axon turning angle quantification, demonstrating Robo-mediated NELL repulsion. Error bars indicate SEM. **G.** Summary matrix indicating presence (closed square) or absence (open ×) of any NELL– Robo binding in each species tested or inferred.

## DISCUSSION

Understanding the evolution and diversification of conserved signaling systems is critical to uncovering how molecular pathways are tuned to support the increasing structural and functional complexity of animal nervous systems. The Robo receptor family provides a compelling model for studying this phenomenon in the context of guidance systems, given its essential role in axon guidance and its expansion and functional divergence across metazoans. Traditionally, Robos are known as receptors for Slit ligands, mediating repulsive axon guidance responses during neural circuit assembly but also fulfilling other Slit-dependent functions in various organismal contexts. However, the more recently discovered mammalian NELL–Robo3 interaction highlights an additional ligand–receptor axis whose evolutionary origins and functional relevance across species had remained poorly understood.

Our results demonstrate that NELL-Robo interactions are an evolutionarily ancient feature of the Robo receptor family, predating vertebrates and emerging at least as early as the last common ancestor of chordates. This revises the current view that NELLs are a recent, mammal-specific innovation co-opted by Robo3. Instead, we show that multiple deuterostome Robos, including those from cephalochordates and cyclostomes, form high-affinity complexes with NELL ligands. These interactions can drive robust repulsive axon guidance, suggesting that NELL–Robo signaling was an ancestral contributor to Robo-mediated nervous system development, potentially complementing or even predating the better-characterized Slit-Robo axis.

Our data, supported by previous work (Pak *et al*, 2020; Yamamoto *et al*, 2019), now suggest that chordate Robos have originated as dual-ligand receptors, with the ability to bind both Slits and NELLs. Mammalian Robo3 appears to have retained NELL responsiveness but lost Slit binding, while Robo1 and Robo2 show only weak or cryptic NELL binding unless their ECD conformation is relaxed. Given that the full-length forms of many chordate Robos appear to mask the NELL-binding site (**Figure 3**), the evolutionary loss of NELL responsiveness may have occurred through conformational changes rather than complete loss of binding domains (except in the case of vertebrate Robo4 (Xu *et al*, 2021)).

From a broader evolutionary perspective, our findings raise the question of whether NELL-Robo interactions were present in the last common ancestor of all bilaterians and subsequently lost in protostomes, or whether this signaling axis arose later within the deuterostome lineage. The absence of NELL genes in many protostome genomes, including *Drosophila* and *C. elegans*, makes it difficult to infer whether loss or *de novo* evolution explains the present distribution. Regardless, the widespread conservation of functional NELL–Robo complexes among vertebrates and basal chordates, together with their biophysical properties, including the tendency to undergo ligand-induced phase separation *in vitro*, suggests that this interaction mode plays a fundamental role in axon guidance.

Although we recovered a NELL-like gene in a sponge, the absence of detectable Robo orthologs in Porifera prevents us from discriminating between two potential scenarios: (1) NELL-Robo coupling arose in a common ancestor of bilaterians and was secondarily lost in protostomes, or (2) the interaction emerged *de novo* in the deuterostome branch.

Together, our phylogenetic, biochemical, biophysical, and functional data reveal that NELL–Robo signaling is not a mammal-specific novelty but a deeply conserved guidance module that emerged at least in the earliest chordates. The ancestral Robo receptor likely recognized both Slit and NELL ligands, with later paralog-specific conformational changes fine-tuning this dual-ligand capacity. However, some extant Robos, e.g. chick Robo3, appear to have retained this ability to interact with both types of ligands, even in the context of the full receptor ECD. Several important questions remain: The discovery that NELL–Robo complexes undergo ligand-triggered phase separation and that these condensates form even in lancelet proteins, may highlight a primordial, physical mechanism for amplifying or spatially restricting repulsive cues at the growth cone through formation of large oligomeric ligand-receptor assemblies, though this needs to be experimentally validated. Further, whether Slit and NELL ligands activate the same downstream effectors or elicit distinct intracellular responses remains unknown. Structural and signaling studies will be necessary to clarify if dual-ligand recognition in ancestral Robos reflected a shared signaling pathway or functionally divergent outputs. It is also unclear whether NELL and Slit can simultaneously bind to a single Robo receptor molecule and, if so, how this affects signaling.

By demonstrating that cephalochordate NELL can drive Robo-dependent axon repulsion in mammalian neurons, we show that downstream signaling machinery for NELLs has remained compatible for ∼500 million years. These findings reposition NELL-Robo interactions as a core component of the axon guidance toolkit and set the stage for future work to dissect how dual-ligand recognition, conformational gating, and biomolecular condensation integrate to sculpt neural circuits across the animal kingdom.

## MATERIALS AND METHODS

We first queried vertebrate NELL1–3 sequences against the NCBI non-redundant, Ensembl, and JGI databases using an iterative, reciprocal BLAST approach. Each candidate hit was then validated by reciprocal BLAST back to human NELLs and inspected for the diagnostic domain architecture (**Figure 1A**). In parallel, we searched the OrthoMCL orthology tables and manually screened predicted proteomes from representative phyla.

### Phylogenetic analysis

Sequences were aligned using MUSCLE (Edgar, 2004), and the maximum-likelihood phylogeny was inferred using RAxML (v8.2.12) (Stamatakis, 2014). The best-fit model of evolution for NELL was WAG + G + I + X (the WAG substitution matrix, gamma-distributed among-site rate variation, invariable sites, and maximum-likelihood equilibrium frequencies). The best-fit model for ROBO was LG + G + I + X. Approximate likelihood ratio statistics were computed using PhyML (v3.0) (Guindon *et al*, 2010) using the model parameters inferred by RAxML. In the maximum-likelihood ROBO3 phylogeny, mammalian sequences were wrongly assigned as the most basal-branching vertebrate lineage, likely because of long-branch attraction caused by their exceptionally high evolutionary rate. We therefore constrained the tree to enforce the correct species topology; all other lineages were congruent with the established species relations.

### Protein expression and purification

All Robo and NELL proteins used in SPR, phase separation and axon guidance assays were expressed using the baculovirus expression system. Either full ECDs or domain truncations were cloned into pAcGP67A (BD Biosciences) with C-terminal hexahistidine tags and co-transfected into Sf9 (*Spodoptera frugiperda*) cells with BestBac2.0 linearized baculovirus DNA (Expression Systems, 91-002) using TransIT-Insect (Mirus, MIR 6104) or Cellfectin II (Thermo Fisher, 10362-100) transfection reagents according to manufacturer’s instructions. Sf9 cells were cultured in SF900 SFM III (ThermoFisher, 12658019) with 2 mM L-glutamine (Cytiva, SH30034.02), 20 µg/ml gentamicin sulfate (Lonza, 17-518L) and 10% fetal bovine serum. Conditioned media from transfected Sf9 cells containing active virus were collected for infection.

For protein expression, High Five cells (*Trichoplusia ni*, BTI-TN-5B1-4) grown in Insect-XPRESS (Lonza, 12-730Q) with 10 µg/ml gentamicin sulfate were infected at 2 × 10^6^ cells/ml density and conditioned media containing secreted protein were harvested 48-66 hrs post-infection. High Five cells were grown in suspension at 27.5°C

Proteins were purified using Ni-NTA agarose resin (Qiagen, 30250) following precipitation of unknown metal chelators by addition of 50 mM Tris pH 8.0, 5 mM CaCl_2_ and 1 mM NiCl_2_ to conditioned media. All proteins were further purified using size-exclusion chromatography (SEC) with Superdex 200 Increase 10/300 (Cytiva) for domain truncation constructs or Superose 6 Increase 10/300 for full-length or complete ectodomain proteins in HEPES-buffered saline (HBS, 10 mM HEPES pH 7.2, 150 mM NaCl).

All protein expression constructs used are tabulated in **Table S1**.

### Extracellular interactome assay (ECIA)

Full-length and truncated versions of proteins were expressed in bait (Fc) and prey (AP) form in *Drosophila* S2 cells as described above. The baits were applied onto protein A-coated 96-well plates washed with PBST, pH 7.5 and incubated overnight at 4C agitating at 500 rpm on an orbital shaker. The bait solutions were discarded and blocking solution (PBS, pH 7.5 with 1% BSA) was applied into the wells followed by 3 hours of incubation at room temperature while agitating on an orbital shaker at 500 rpm. Protein expression is confirmed by western blotting using iFluor 488-coupled anti-His antibodies (Genscript, catalog no. A01800).

### Surface plasmon resonance (SPR)

Proteins expressed in High Five cells as described above were biotinylated using Sulfo-NHS-Biotin in 0.1 M MES pH 6.4 to allow only for only the monobiotinylation of the N-terminal amine groups. Proteins were purified from biotin excess using SEC on Superdex 200 or Superose 6 10/3000 columns. SPR experiments were performed on a Biacore 8K instrument (Cytiva). Biotinylated proteins were immobilized on streptavidin-coupled SPR chips (Cytiva). Analytes were diluted in HBS with 0.05% Tween-20 and 0.1% BSA and flown over the ligands in the same buffer at 30 µl/min. The results were analyzed using the BIAevaluation software (Cytiva), and steady-state data were fit to one-site binding model with a linear non-specific binding term in Prism (version 11.0.2).

### Phase separation assay

For initial observational studies and turbidity assays, Robo and NELL proteins expressed in High Five cells as described above were mixed at concentrations above 1 µM in equal volumes and allowed to rest at room temperature for at least 5 min. Turbidity assays were performed 5 min after mixing by measuring absorbance at 600 nm for each solution. For western blots, 1000 µl mixed solutions were allowed to settle overnight and were centrifuged at 13000 × g for 10 minutes. The supernatant was set aside and the pellet was mechanically resuspended in 1000 µl HBS. Both supernatant and resuspended pellet were mixed with 6x SDS sample loading buffer and boiled at 95°C for 10 min before loading 50 µl onto an SDS-PAGE gel. The gel was stained with Coomassie Blue for 1 hr and destained with 10% acetic acid and 50% ethanol. The gel was imaged in a BioRad Chemidoc MP system.

For fluorescence microscopy, human Robo3 was labeled with NHS-fluorescein (ThermoFisher 46409) and human NELL2 was labeled with NHS-Cy5 (Millipore Sigma ATEH9B9AC747). Fluorophores were added in a 10:1 ratio of fluorophore:protein for the labeling reaction buffered with cold 50 mM MES pH 6.5. Reactions were incubated for 1 hr on ice then quenched with 100 mM Tris pH 8.0. Excess fluorophore was removed using SEC. Images were taken on an Olympus “live cell” DSU spinning disk confocal.

### Animals

All animal procedures were approved by the Brown University Institutional Animal Care and Use Committee and performed in accordance with National Institutes of Health guidelines. Mice carrying the *Robo3* null allele have been described before and were genotyped as previously reported (Sabatier *et al*, 2004). *Robo3* knockout embryos were generated from heterozygous *Robo3^+/−^* intercrosses. Embryonic day (E) 0.5 was defined as the day a vaginal plug was detected. Both male and female embryos were included in the study.

### Spinal cord electroporation and primary neuron culture

Spinal cord electroporation, tissue dissociation, and neuronal cell culture were carried out essentially as described before (Pak *et al*, 2020). In short, different Robo expression constructs, together with an RFP expression vector (at 3:1 molar ratio, combined DNA concentration of 100 ng/µl; for the “no rescue” condition, Robo expression construct was omitted), were injected into the spinal cord central canal of E11.5 *Robo3^−/−^*embryos, followed by bilateral electroporation. All expression constructs are listed in Table S1. After electroporation, the dorsal half of the cervical and thoracic spinal cord was microdissected and dissociated into single cells, and neurons were cultured on glass coverslips coated with 100 µg/ml Poly-D-Lysine (Sigma) and 5 μg/ml mouse N-Cadherin (R&D Systems). Neurons were used for Dunn chamber axon turning assays 16–24 h after plating.

### Dunn chamber axon turning assay

Axon turning assays were performed as previously described (Pak *et al*, 2020). In short, pre-cultured neurons were exposed to a gradient of NELL protein in a Dunn chamber and live-imaged over the course of 2 h. To establish gradients, 50 ng/ml of NELL protein was added to the outer well of the Dunn chamber. For mock rescue and rescue with mouse Robo3.1, neurons were exposed to mouse NELL2 (R&D Systems); for rescue with lancelet Robo or zebrafish Robo1/Robo3, neurons were exposed to purified recombinant lancelet NELL or zebrafish NELL2b (see protein expression and purification), respectively. After live imaging, neurons were stained with antibodies against TAG-1/Cntn2 to distinguish commissural from ipsilaterally projecting neurons and for RFP to identify electroporated neurons, as reported before (Pak *et al*, 2020).

### Quantification of axon turning

Quantification of axon turning angles was carried out as previously described (Pak *et al*, 2020). Turning angles from electroporated commissural and ipsilaterally projecting *Robo3^−/−^* neurons were pooled, as both types of neurons become NELL-responsive when forced to express mouse Robo3.1 (Pak *et al*, 2020) or any of the lancelet and zebrafish Robos (**Figure S3A)**. A minimum of 18 neurons (and up to 70 neurons) in each Dunn chamber replicate, for 3-4 biological replicates per experimental condition (tissue from each embryo receiving a specific rescue construct counting as a single replicate), were analyzed. Turning angles were averaged across neurons for each replicate, and the means across multiple replicates were analyzed for statistical significance using a one-way ANOVA with Dunnett’s *post hoc* test (n and *p* are indicated in the figures). SEC profiles and SDS-PAGE gels for purified NELL proteins used in the assay are included in **Figures S3B,C**.

## Acknowledgments

We thank Hyun Lee and the Biophysics Core Facility at the University of Illinois Chicago for technical assistance.

## Author contributions

E.Ö., A.J. and J.S.P. designed the project. J.S.P, W.I.N., R.K. performed biochemistry experiments. L.P. performed the neurobiology experiments. Y.P. performed the phylogenetic analysis. A.J., E.Ö and J.W.T. provided supervision and acquired funding. E.Ö., A.J., W.I.N., L.P., J.S.P., and Y.P. wrote the manuscript.

## Data and materials availability

All data needed to evaluate the conclusions in the paper are present in the paper or the Supplementary Materials. The plasmids are available from E.Ö. and A.J.

---

**Figure S1.**
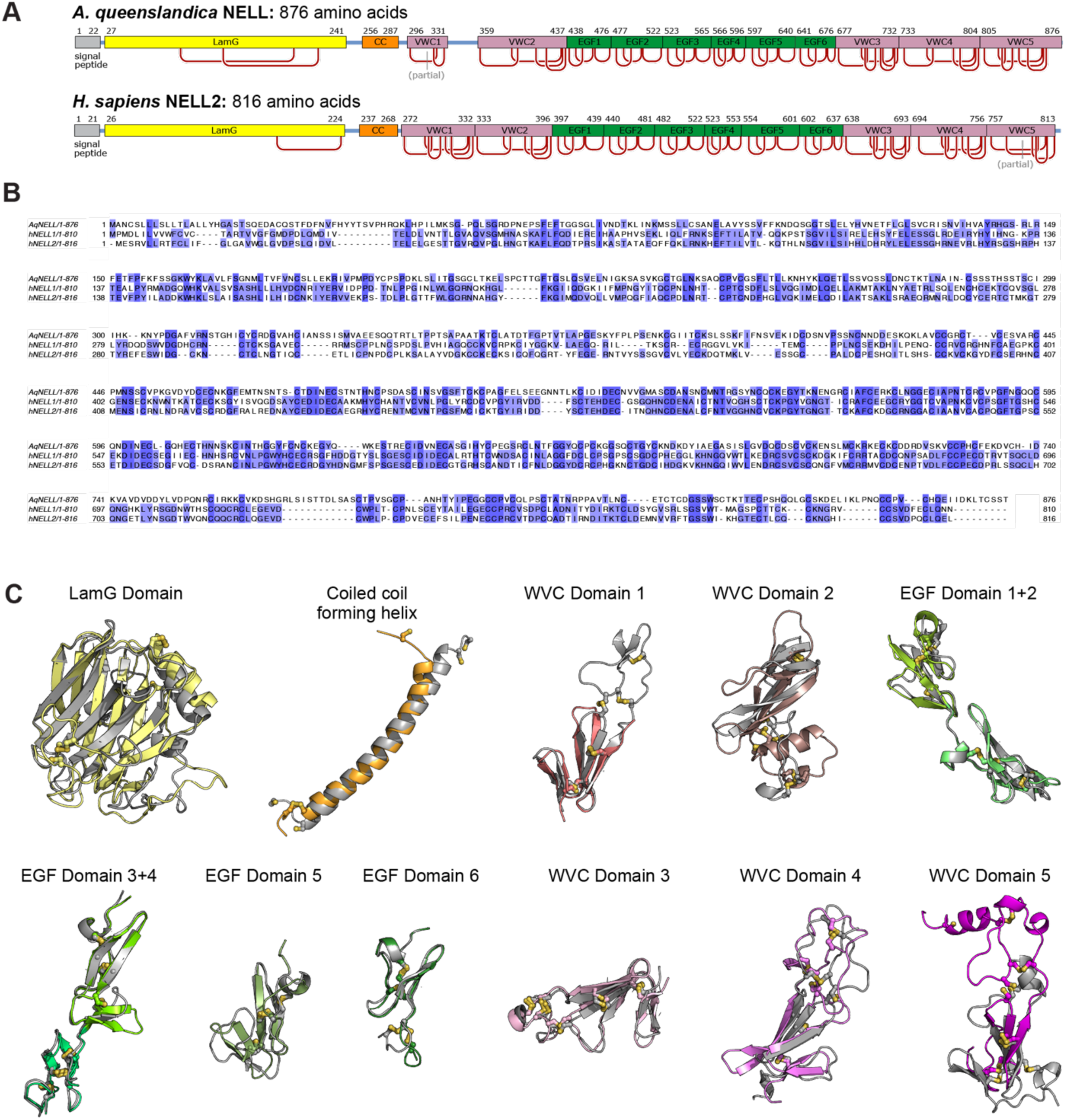
Conserved sequence and structural features of poriferan and human NELLs. **A.** Domain architectures of *A. queenslandica* NELL (XP_003383458.2) and human NELL2 as inferred from Alphafold predictions. **B.** Sequence alignment for *A. queenslandica* (Aq) and human (h) NELLs. Sequence identity between AqNELL and hNELL1 or hNELL2 are both 26%. **C.** Structural alignments of domains of *A. queenslandica* NELL and human NELL2. Alphafold model for human NELL2 is in gray, while A. queenslandica NELL are in various other colors.

**Figure S2.**
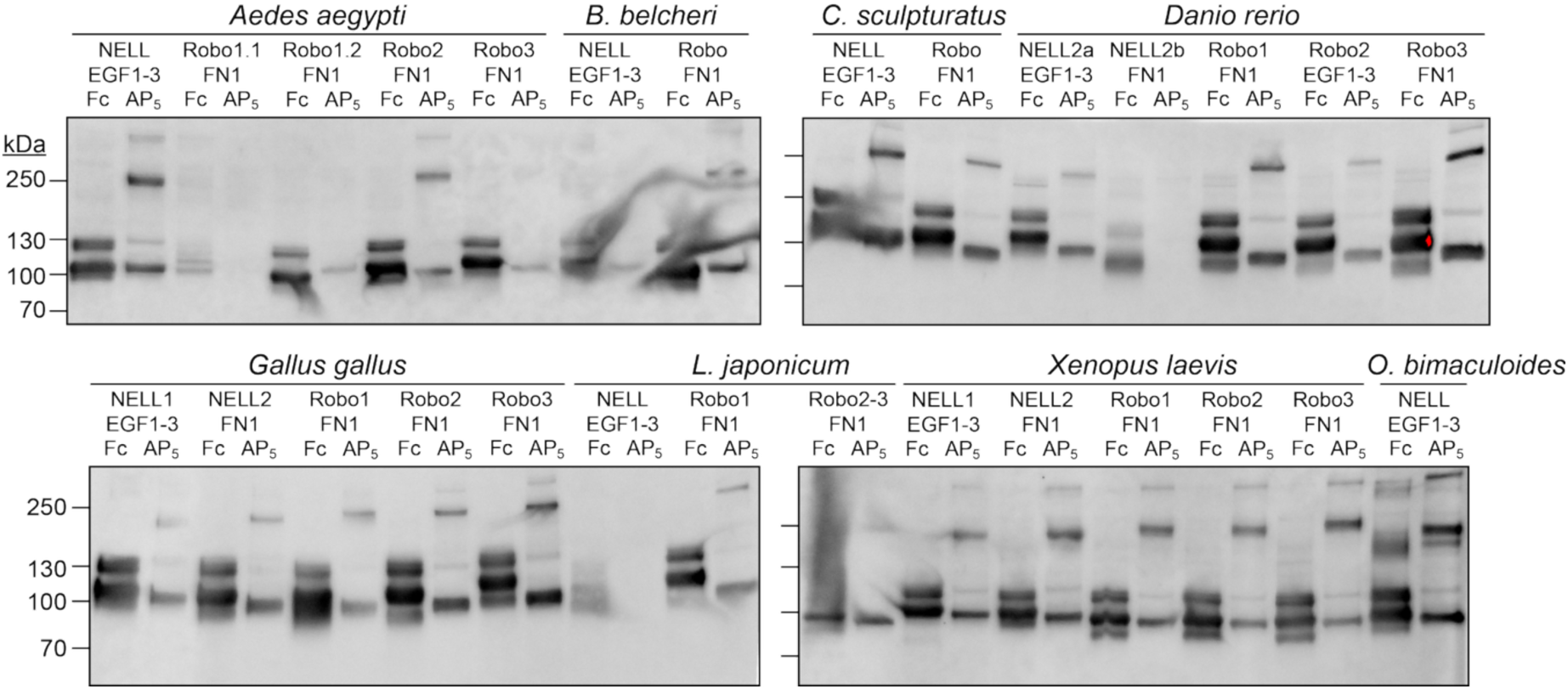
Expression of Robo and NELL constructs used in ECIA. Anti-His tag western blots of proteins secreted from *Drosophila* S2 cells for ECIA (Fig. 2), using a non-reducing protein loading buffer. The Fc and AP_5_ constructs are disulfide-mediated dimers and pentamers, but intermediate oligomeric states are often observed on gels, resulting in multiple bands for each construct. Not accounting for heterogeneity and added mass of N-linked glycans (+5-15%), expected approximate sizes are EGF1-3-Fc: 48 kDa × 2; EGF1-3-AP_5_: 81 kDa × 2; FN1-Fc: 47 kDa × 2; FN1-AP_5_: 80 kDa × 2.

**Figure S3.**
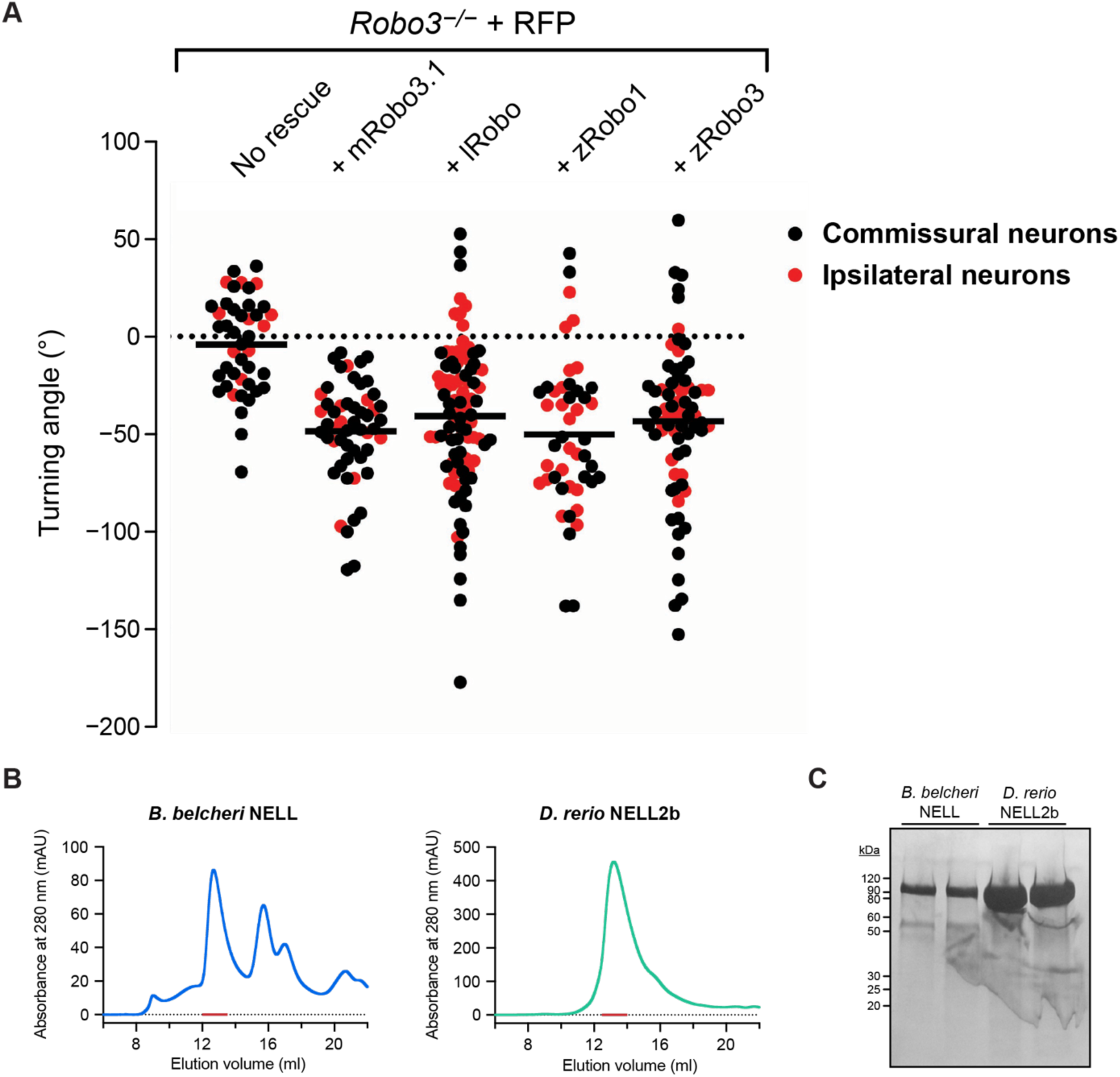
Expression and functional quantification for Dunn-chamber assays. **A.** Distribution of individual axon turning angles for commissural and ipsilaterally projecting neurons (m = mouse, l = lancelet, z = zebrafish). **B.** Size-exclusion chromatograms of *B. belcheri* NELL and *D. rerio* NELL2b preparations used in the Dunn-chamber assays. Red bars show the fractions collected and used. A Superose 6 10/300 column (Cytiva) was used, with elutions volumes consistent with trimeric NELL molecules. Absorbance path length: 0.2 cm. **C.** Corresponding SDS-PAGE gels showing the purity of the NELL samples used in the Dunn-chamber assays.

**Table S1.**
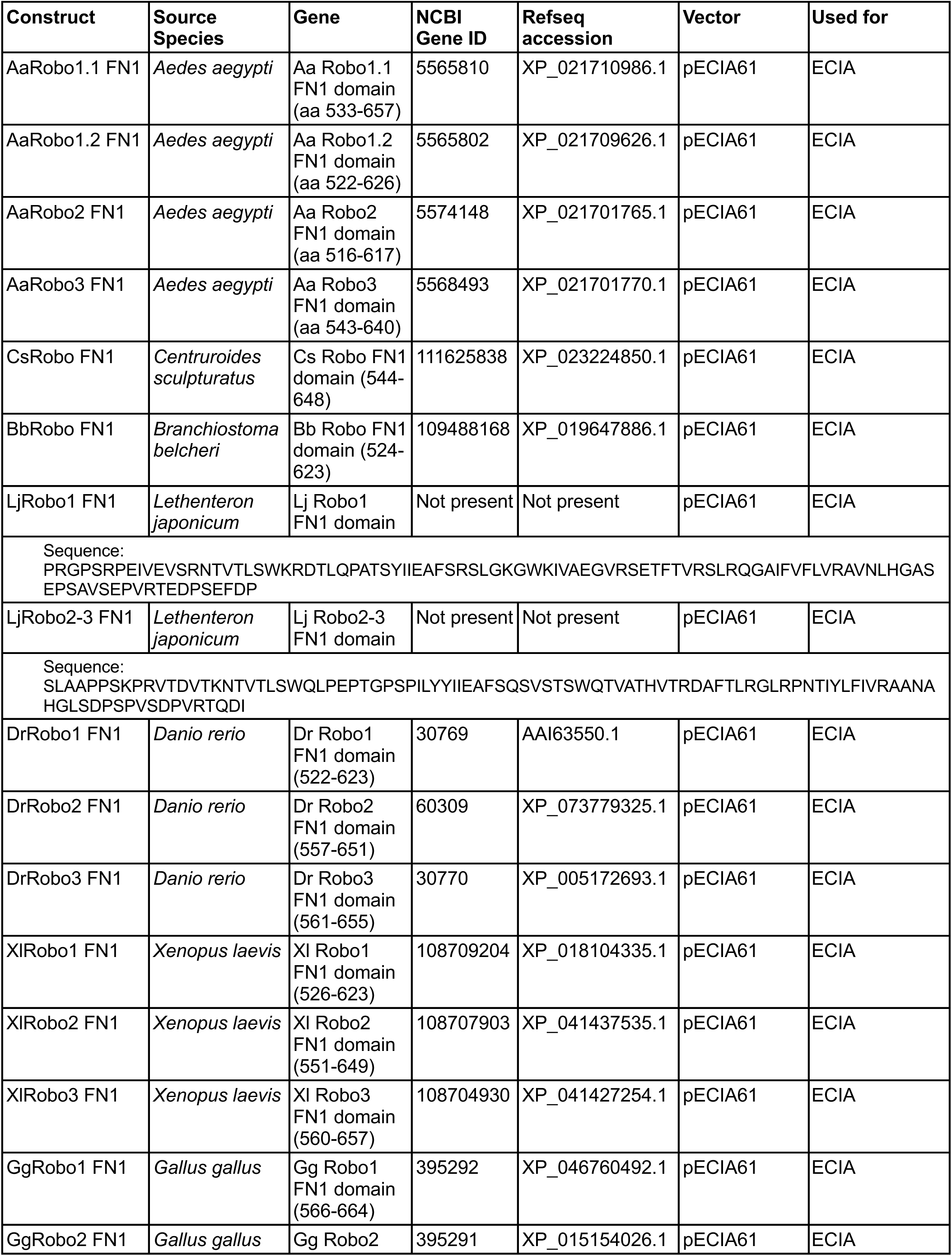
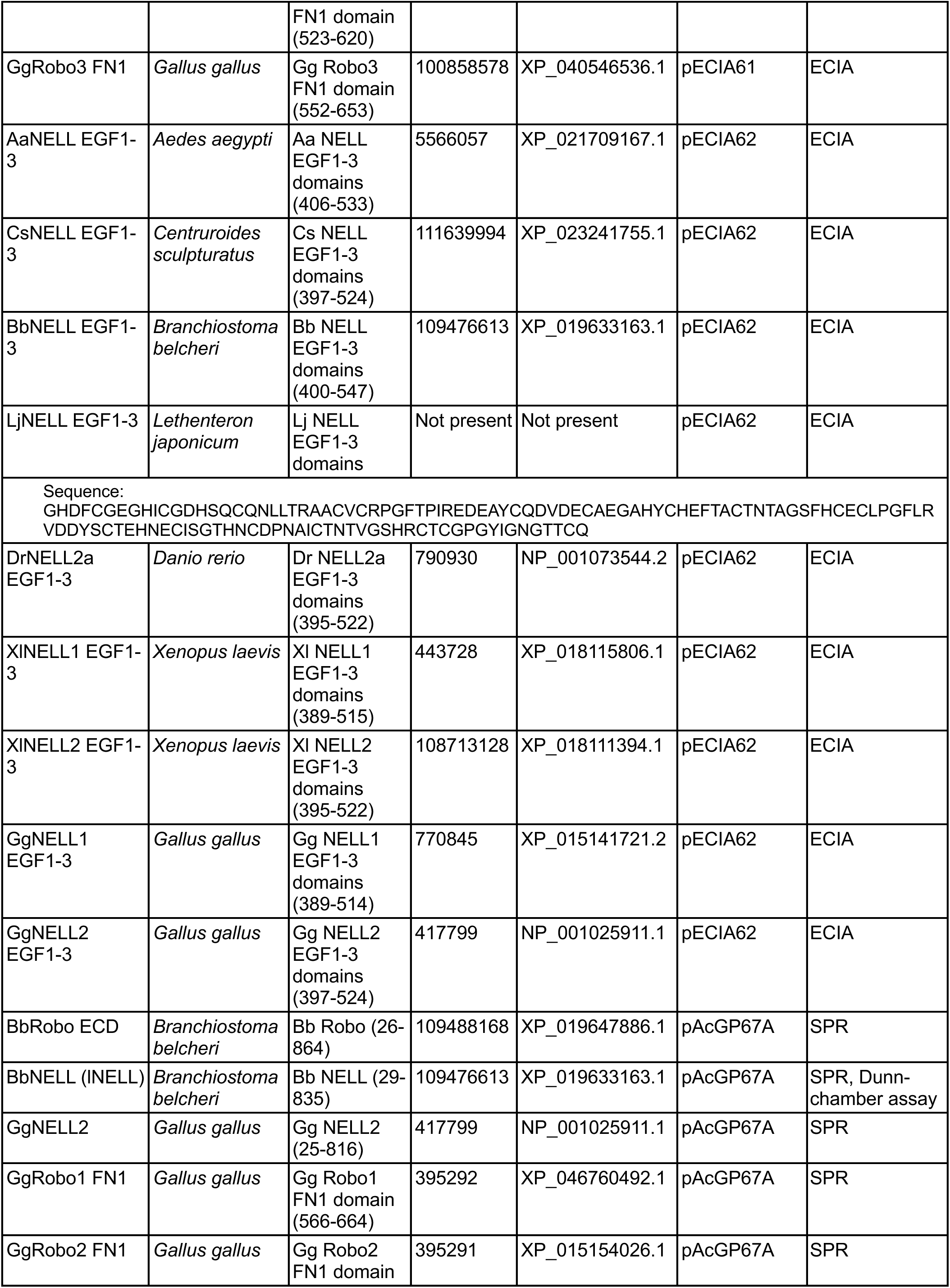
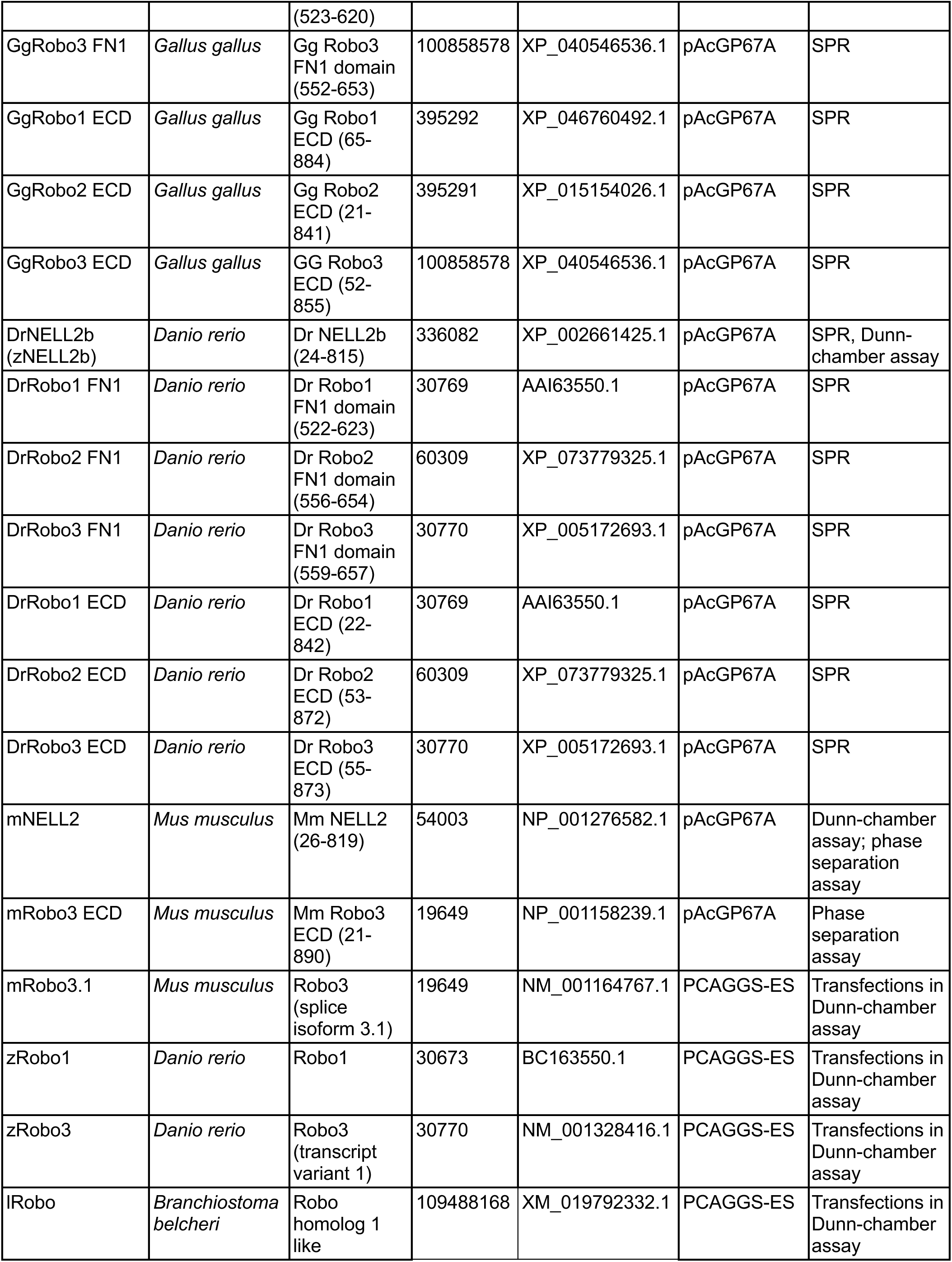
Plasmids created for and used in this study.

